# Enhanced restoration of visual code after targeting on bipolar cells compared to retinal ganglion cells with optogenetic therapy

**DOI:** 10.1101/2024.07.22.604613

**Authors:** Jessica Rodgers, Steven Hughes, Aghileh Ebrahimi, Annette E Allen, Riccardo Storchi, Moritz Lindner, Stuart N Peirson, Tudor Badea, Mark W Hankins, Robert J Lucas

**Author notes:** Contributed equally (Jessica Rodgers, Steven Hughes).

## Abstract

Optogenetic therapy is a promising vision restoration method where light sensitive opsins are introduced to the surviving inner retina following photoreceptor degeneration. The cell type targeted for opsin expression will likely influence the quality of restored vision. However, a like-for-like pre-clinical comparison of visual responses evoked following equivalent opsin expression in the two major targets, ON bipolar (ON BCs) or retinal ganglion cells (RGCs), is absent. We address this deficit by comparing stimulus-response characteristics at single unit resolution in retina and dorsal lateral geniculate nucleus (dLGN) of retinally degenerate mice genetically engineered to express the opsin ReaChR in *Grm6-* or *Brn3c*-expressing cells (ON BC vs RGCs respectively). For both targeting strategies, we find ReaChR-evoked responses have equivalent sensitivity and can encode contrast across different background irradiances. Compared to ON BCs, targeting RGCs decreased response reproducibility and resulted in more stereotyped responses with reduced diversity in response polarity, contrast sensitivity and temporal frequency tuning. Recording ReaChR-driven responses in visually intact retinas confirmed that RGC-targeted ReaChR expression disrupts visual feature selectivity of individual RGCs. Our data show that while both approaches restore visual responses with impressive fidelity, ON BC targeting produces a richer visual code better approaching that of wildtype mice.

## INTRODUCTION

Optogenetic therapy is a promising vision restoration approach for retinal degeneration. In retinas without rod and cone photoreceptors, the missing visual input can be replaced by genetically engineering surviving retinal cells to express light-sensitive proteins called opsins^1^. This approach is appropriate for patients suffering photoreceptor loss regardless of underlying cause and has been validated in numerous preclinical studies employing animal models of retinal degeneration ^2,3^. Several clinical trials are underway, with an early report of successful, albeit limited, vision restoration ^4^.

An important decision point for any optogenetic intervention is which cell type(s) in the retina to target for photopigment expression. The suitability of AII amacrine cells ^5^ and surviving cones ^6^ have been explored, but most work on this topic has focused on either ON bipolar cells (ON BCs ^7–13^) or retinal ganglion cells (RGCs ^14–21^). These two cell types offer distinct advantages and limitations. In favour of RGCs is ease of transduction and resilience in the face of progressive degeneration. RGCs are the main population transduced following intravitreal injection using ubiquitous promoters, such as CAG or CMV, with available adeno-associated virus serotypes ^22^. In comparison, bipolar cells are more challenging to transduce and require a combination of cell-specific promotors ^8,23–25^ and modified capsids ^26–28^. RGCs are also less affected by the retinal reorganisation that accompanies progressive degeneration, while BCs are more subject to cell death and exhibit greater morphological and genetic changes after retinal degeneration than RGCs ^29–31^. In principle this makes RGCs more reliable hosts for optogenetic actuators.

The major argument in favour of ON BCs is that targeting these cells may better recreate the natural visual code. Here, retinal circuitry upstream of RGCs performs important computations allowing diversity in visual feature selectivity in the retinal output ^32,33^. Introducing optogenetic actuators to RGCs runs the risk of replacing this complex visual code with a stereotyped visual response that fails to recreate the expected visual feature selectivity for many ganglion cell types. By contrast, optogenetic signals originating in ON BCs would propagate through inner-retinal circuits allowing the possibility that they will be subject to many of the same computations as native photoreceptor-derived inputs. The most obvious example of such processing is the generation of separate ON and OFF representations of the visual scene and, indeed, ON BC optogenetic interventions can recreate such diversity in the visual code ^8,11,13,34^.

Understanding how the visual code is impacted by choosing to target optogenetic actuators to ON BCs vs RGCs is thus an important step in optimising this therapy. Comparisons of visual response following expression of the same opsin targeted to different cell types have asked whether ON BC targeting strategies offer advantages over approaches biased towards RGC expression ^23,25,35^. Based upon population level analyses of light-evoked activity in RGCs these have reported that ON BC targeting can reduce photosensitivity ^23,25^ (although see Linder et al. ^35^) and alter temporal frequency tuning ^35^. However, those studies fall short of a detailed description of the visual code at single unit resolution as required to reveal its complexity. Moreover, they employed viral gene delivery to achieve opsin expression, introducing unavoidable variability in the amount of opsin expressed across cells and allowing the possibility that any differences in response properties arise from the difference in viral transduction efficiency between cell types.

A clearer picture of the relative advantages of ON BC vs RGC targeting thus awaits a quantitative comparison of the visual code at single unit resolution without the variability inherent in viral gene delivery. We have previously used a transgenic mouse line to achieve more uniform and pan-retinal expression of the optogenetic actuator ReaChR across ON BCs and shown that this strategy has an impressive ability to recreate features of the wildtype visual code when applied to the *Pde6^rd1^* model of retinal degeneration ^34^. Here, we adopted a similar approach to producing ReaChR expression across a population of RGCs in order to facilitate a like-for-like comparison of visual restoration by the same opsin expressed under the same promoter in the two different cell types. We find that both ON BCs and RGCs are suitable targets for high fidelity visual responses, but that targeting ON BCs does indeed better recreate the diversity of the natural visual code.

## RESULTS

### Transgenic mouse models of ON bipolar cell vs retinal ganglion cell-targeted optogenetic vision restoration

To achieve more standardised opsin expression than is achievable with viral gene delivery, we generated versions of the *Pde6b^rd1^* model of retinal degeneration ^36,37^ bearing a floxed ReaChR-mCitrine cassette in the Rosa26 locus ^38^, combined with a transgene providing Cre recombinase expression under either Grm6 (termed here ReaChR Grm6 rd/rd ^34,39^) or Brn3C (termed here ReaChR Brn3cr rd/rd ^40^) promoters. This results in retinally degenerate mice with expression of the red-shifted channelrhodopsin, ReaChR (a light-sensitive ion channel ^41^) in ON BCs and a subset of RGCs, respectively. Immunohistochemical analysis of retinal sections from 5 month old animals confirmed the expected degeneration of the photoreceptor cell layer and revealed ReaChR expression in cells at the outer portion of the inner nuclear layer in Grm6cre, and in the ganglion cell layer in Brn3cre mice, consistent with expression in ON BCs and RGCs (**Fig 1A**). Dendrite stratification from Brn3c-expressing RGCs was found in both the on and off sublaminae of the inner plexiform layer, consistent with previous reports that this Cre-driver line targets monostratified ON and OFF, as well as bistratified, RGC subtypes^40^.

**Figure 1.**
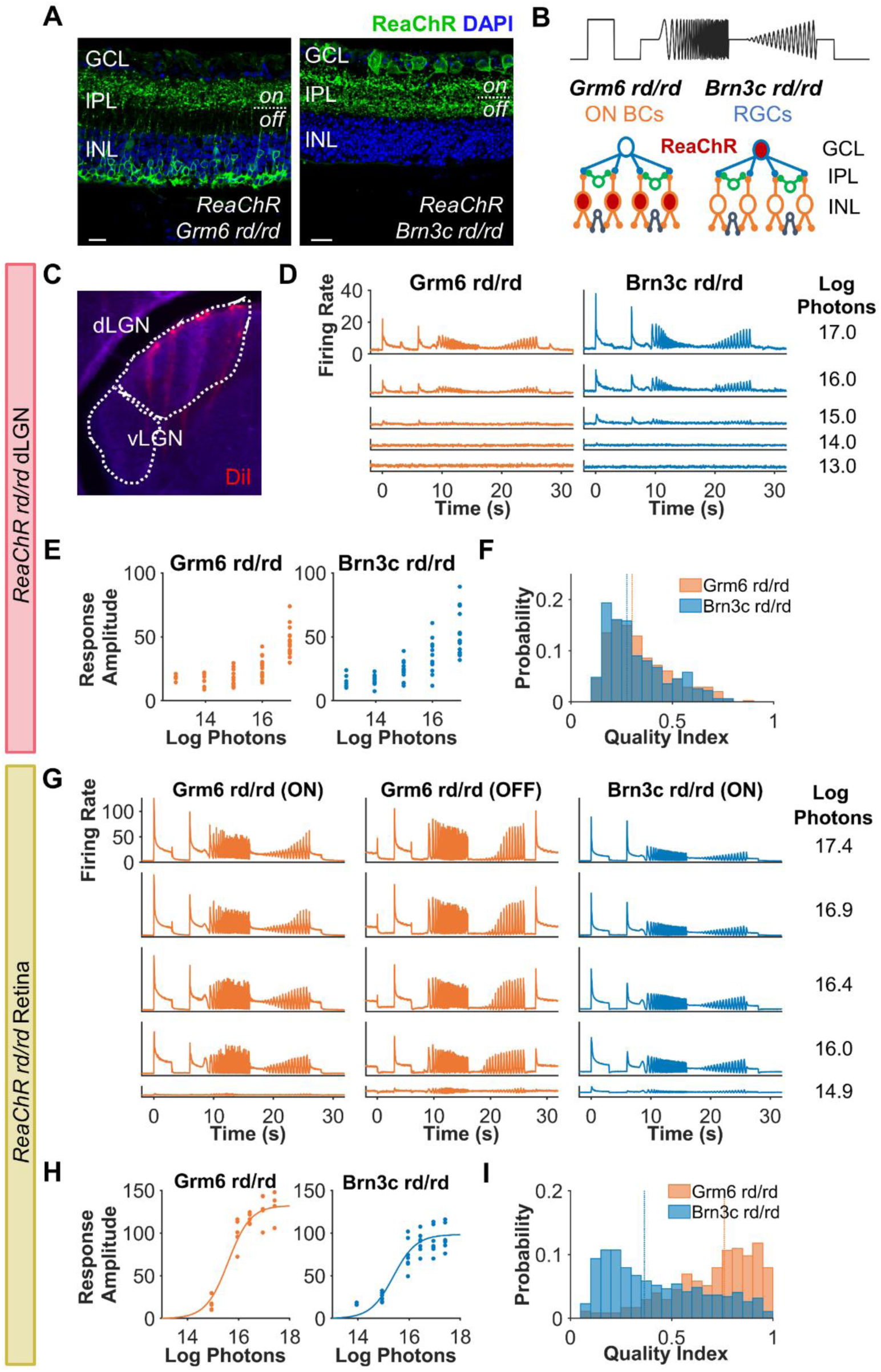
a) Immunohistochemistry from retinal sections from ReaChR Brn3C rd/rd (Brn3c) and ReaChR Grm6 rd/rd (Grm6) stained for DAPI (blue) and ReaChR-mCitrine (GFP). Scale bar = 20µm. GCL = ganglion cell layer, IPL = inner plexiform layer, INL = inner nuclear layer, on = ON sublamina of IPL, off = OFF sublamina of IPL. b) Chirp stimulus (top) and schematic of experiment. ReaChR was expressed in a subset of RGCs or ON bipolar cells (ON BCs) and visual responses recorded from RGCs and dLGN neurons using multi-electrode arrays. c) LGN targeting using multi-electrode arrays. Location of shanks in dLGN during *in vivo* electrophysiology recordings are shown by fluorescent labelling using DiI (red). d,g) Mean firing rate for light responsive (LR) c) dLGN units (n = 310 for Grm6 and 348 for Brn3c) & e) LR retinal units (n = 343 for Grm6 ON, n = 116 for Grm6 OFF, n = 880 for Brn3c) to chirp stimulus across intensities. e,h) Mean maximum baseline-subtracted firing rate to step stimulus for LR units from d) each dLGN placement (N = 16 for Grm6 and 14 for Brn3c) and f) each retinal recording (N = 5 for Grm6 and N = 7 for Brn3c) across intensities. Data in f) are fit with irradiance response curve. f,i) Histogram of quality index for LR units at brightest intensity tested from e) dLGN units at 16.97 log photons (n = 310 Grm6, n = 348 for Brn3c) and f) retina units at 17.4 log photons (n = 774 Grm6 units, n = 827 Brn3c units)

### Population level-responses – amplitude, sensitivity and sensitivity normalisation

We examined responses to a full-field chirp stimulus, previously used to characterise visual response diversity in mice ^32^. The chirp comprised a 3s light step from dark followed by sinusoidal modulations of increasing temporal frequency or contrast (**Fig 1B**) and was presented at a range of irradiances. To assess visual responses, we first set out to record visually evoked responses in the dorsal lateral geniculate nucleus (dLGN) in urethane-anesthetised mice (**Fig 1C**). At 5 months of age mice homozygous for the *Pde6b^rd1^* mutation (termed here rd/rd) have advanced retinal degeneration and dLGN visual responses are restricted to very sluggish and low sensitivity changes in firing driven by the low acuity, inner retinal, photoreceptor melanopsin ^42^. As a first assessment of the success of ReaChR expression in either ON BCs or RGCs at restoring visual responses we reviewed mean firing rate profiles of spike sorted single units from the dLGN of Grm6 or Brn3c mice across repeats of the chirp. In both genotypes, high amplitude modulations in mean firing rate associated with the stimulus were apparent at higher irradiances (N = 16 electrode placements from 7 mice for ReaChR Grm6 rd/rd and N = 14 placements from 6 mice for ReaChR Brn3c rd/rd, **Fig 1D**). We then quantified response amplitude as the peak change in baseline-subtracted firing rate during or just after the 3s step across an epoch designed to capture both ON and OFF excitation responses. This parameter was positively correlated with irradiance in both genotypes, with no detectable difference in response to the brightest light (**Fig 1E**, median = 45 spikes/s for ReaChR Grm6 rd/rd and 47 spikes/s for ReaChR Brn3c rd/rd, U = -0.68, p = 0.493, Mann-Whitney U-test used for statistical comparisons between these genotypes unless otherwise specified). As a more comprehensive assessment of response calibre, we calculated a quality index describing response reproducibility (from 0 to 1, where 1 is identical response to all trials^32^). Quality index was slightly higher in the ON BC compared to RGC targeted ReaChR, but showed substantial variation across units in both genotypes (Median QI = 0.3 for ReaChR Grm6 rd/rd, 0.26 for ReaChR Brn3c rd/rd, U = 4.187, p < 0.001, **Fig 1F**).

Having recorded the thalamic response to visual stimulation, we next turned to a direct assessment of the retinal output by recording extracellular activity across the ganglion cell layer (GCL) of retinal explants using multielectrode arrays. As in the dLGN, the chirp stimulus induced notable modulations in firing rate across the population of GCL neurons, especially at higher irradiances (**Fig 1G**; plotted separately for units excited vs inhibited by the light step, termed ON and OFF respectively, in ReaChR Grm6 rd/rd following Rodgers et al. ^34^. Few OFF units were found for ReaChR Brn3c rd/rd, see below, so only ON units are shown). To estimate sensitivity, we constructed an irradiance response curve (IRC) based on change in baseline-subtracted firing rate for each retinal recording (N = 5 retinas from 5 ReaChR Grm6 rd/rd mice and N = 7 retinas from 4 ReaChR Brn3c rd/rd mice). There was no substantial difference in photosensitivity (Log EC50, intensity that produced half maximum amplitude response, was 15.6 for ReaChR Grm6 rd/rd and 15.4 photons/cm^2^/s for ReaChR Brn3c rd/rd, **Fig 1H**), although the response amplitude at brightest intensity tested was attenuated for ReaChR Brn3c rd/rd compared to ReaChR Grm6 rd/rd (105.9 and 138 spikes/s respectively, U = 2, p = 0.010). Enhanced response amplitude in ReaChR Grm6 rd/rd was also apparent in the quality index, which was markedly skewed to higher values in this genotype compared to ReaChR Brn3C rd/rd (median QI = 0.74 for Grm6, 0.4 for Brn3c, U = 19.24, p < 0.001, **Fig 1I**).

The vertebrate visual system adjusts its sensitivity according to background light intensity in order to encode visual contrast across large differences in ambient illumination. Though ReaChR-derived vision, irrespective of whether expressed in ON BCs or RGCs, has a high absolute threshold, it is still important to determine if ReaChR-driven responses show sensitivity normalisation at higher light levels, as the alternative would be saturation. We therefore compared responses to contrast modulations at two irradiances within the ReaChR sensitivity range. At the higher mean irradiance (light grey line in **Fig 2A)**, most elements of the contrast chirp stimulus lie above the saturating irradiance for responses to the simple light step (red line in **Fig 2A**). Conversely, all elements of the dimmer chirp (black line in **Fig 2A**) lay within the ReaChR dynamic range as defined by the step response. Plots of mean firing rate across contrast curves revealed that both genotypes showed high amplitude modulations in firing at both irradiances (**Fig 2C**; shown separately for ON and OFF units in ReaChR Grm6cre rd/rd). Moreover, contrast response relationships confirmed that both genotypes were able to track a wide range of contrasts at both backgrounds (**Fig 2D)**. The implication of sensitivity normalisation is supported by similarities in the contrast level that produced half maximum response amplitude (C50) in the face of the 10x difference in mean irradiance (C50 for 16.6 vs 15.6log photons/cm^2^/s = 51% and 46% for ReaChR Grm6 rd/rd ON, 49% and 43% for ReaChR Grm6 rd/rd OFF, 73% and 76% for ReaChR Brn3c rd/rd ON, respectively).

**Figure 2.**
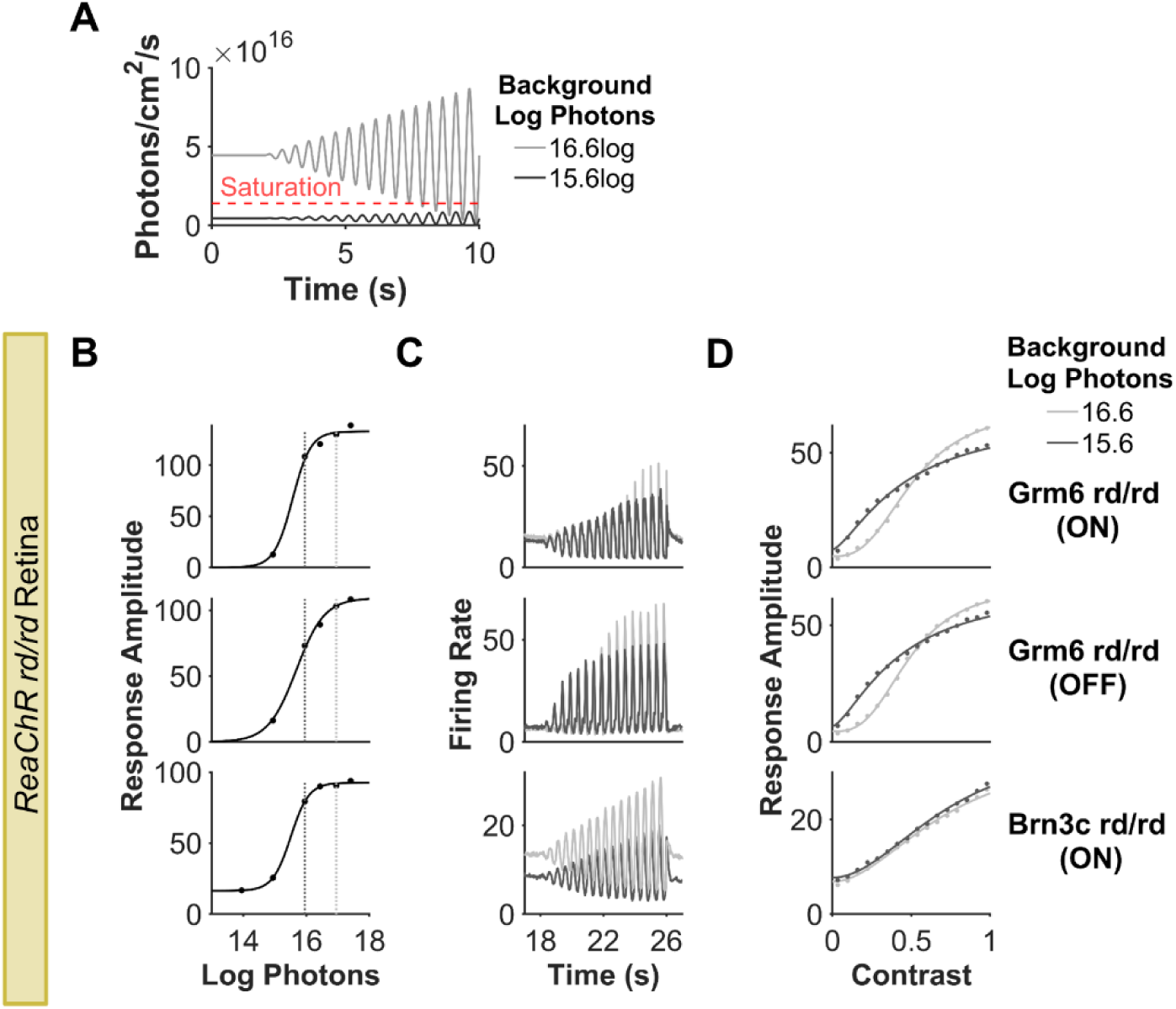
a) Intensity of contrast chirp stimulus at two background irradiances (16.6 and 15.6log photons/cm^2^/s). For responses saturating at 16log (dashed red line), contrast modulations cannot be tracked without sensitivity normalisation. b-d) Comparison of contrast coding at two different background irradiances, b) marked by vertical dashed lines on irradiance response curves c) Firing rate to contrast chirp stimulus and d) contrast sensitivity curves at 15.6 log and 16.6 log photons background. Data in b-d) are from n = 343 for ReaChR Grm6 rd/rd ON, n = 116 for ReaChR Grm6 rd/rd Grm6 OFF, n = 880 for ReaChR Brn3c rd/rd retinal units. Units are divided into ON and OFF as described in Rodgers et al. (2023) to aid visualisation of contrast chirp responses.

### RGC targeting reduces variability in response polarity and kinetics

Having established that RGC targeting using ReaChR Brn3c rd/rd mouse supported robust visual responses with equivalent sensitivity and sensitivity normalisation to that of ReaChR Grm6 rd/rd, we turned to question of whether the identity of the target cells would impact the visual code. Response latency in the retina was reduced in the ReaChR Brn3c rd/rd compared to ReaChR Grm6 rd/rd (**Fig 3A**, median latency for ON response to step = 100ms for Grm6, 75ms for Brn3c, U = 9.98, p < 0.001 at sub-saturating intensity 15.95 log photons/cm2/s), consistent with introduction of ReaChR later in the visual pathway. A survey of responses to step stimulus at single unit resolution (**Fig 3B-C**) indicated a bias towards units meeting an objective classification of ‘ON’ excitation in ReaChR Brn3c rd/rd relative to ReaChR Grm6 rd/rd retinas. To quantify the magnitude of this realignment we calculated an ON-OFF bias index for each unit (from -1 = OFF to 1 = ON; Fig 3C), which confirmed a significant shift towards ON responses in ReaChR Brn3c rd/rd relative to ReaChR Grm6 rd/rd (**Fig 3D**, median = 0.81 and 0.4, respectively, U = -15.57, p < 0.001). Indeed, units with OFF or ON/OFF responses were almost completely lacking from ReaChR Brn3c rd/rd retinas. We further quantified step responses in terms of their persistence. A transience index (from 0 – highly transient to 1 – highly sustained) revealed diversity in both genotypes (**Fig 3E**), but a significant bias towards more sustained responses in ReaChR Brn3c rd/rd (median = 0.25 for Grm6, 0.36 for Brn3c, U = -11.19, p < 0.001).

**Figure 3.**
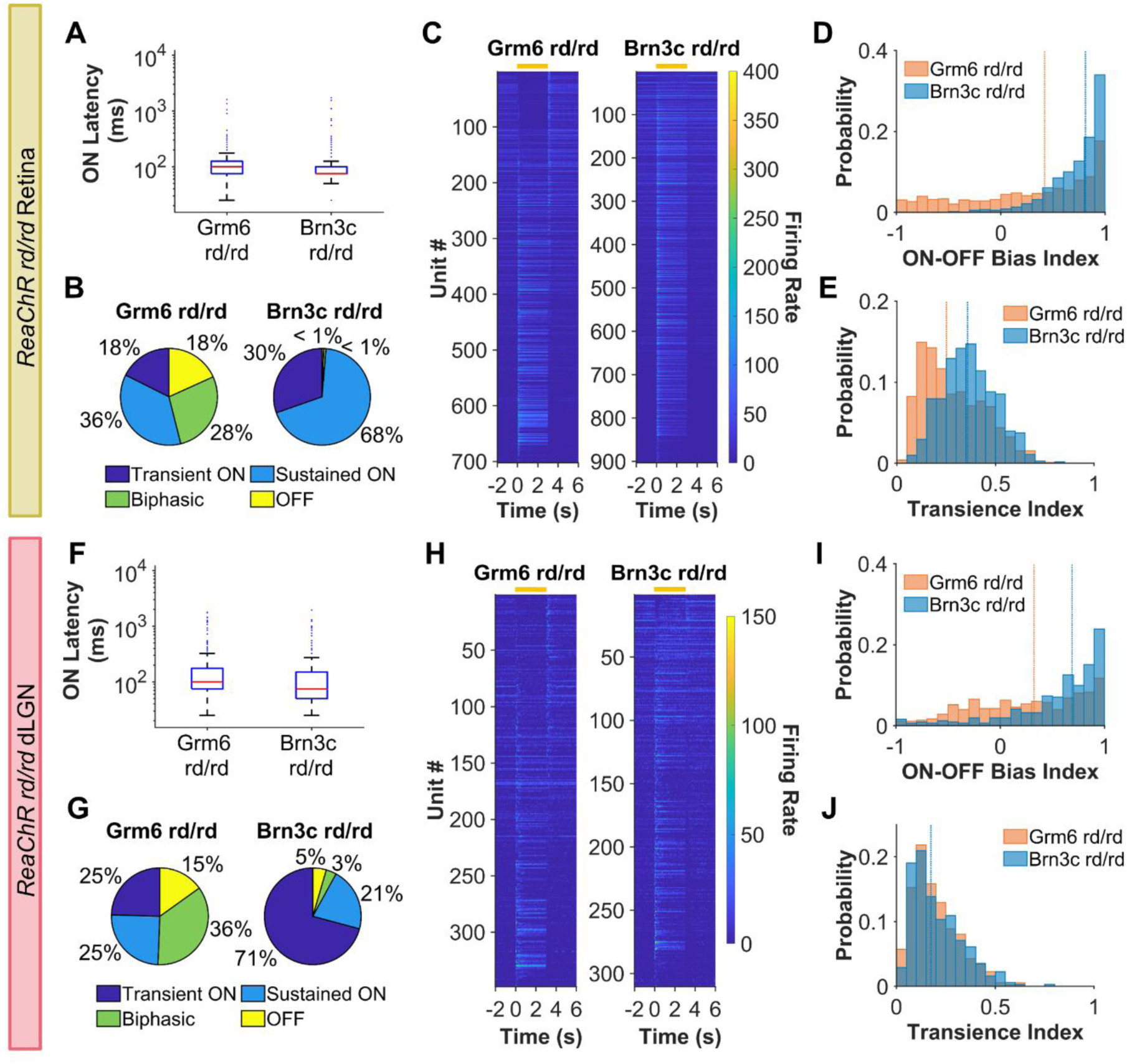
a,f) Latency to onset of step for a) retinal units (N = 662 for ReaChR Grm6 rd/rd and 896 for ReaChR Brn3c rd/rd) and f) dLGN units (N = 283 for ReaChR Grm6 rd/rd and 232 for ReaChR Brn3c rd/rd) b,g) Response type classification c,h) Heatmap of mean PSTH for step stimulus (ON from 0-3s) for LR units ordered by bias index from OFF (top) to ON (high). Each row represents an individual unit, yellow bar shows timing of step stimulus d,i) ON-OFF bias index e,j) Transience index Data in b-e) are from retinal units (N = 702 for ReaChR Grm6 rd/rd and 903 for ReaChR Brn3c rd/rd) at and in f-h) from dLGN (N = 348 for ReaChR Grm6 rd/rd and 310 for ReaChR Brn3c rd/rd).

Some differences in step responses between genotypes were apparent also in the dLGN. Just as in retina, response latency at sub-saturating irradiance (16.97 log photons/cm^2^/s) was reduced in ReaChR Brn3c rd/rd compared to ReaChR Grm6 rd/rd dLGN (**Fig 3F**, median = 100ms for Grm6, 80ms for Brn3C, U = 5.44, p < 0.001). There remained significantly more variability in polarity in ReaChR Grm6 rd/rd compared to ReaChR Brn3c rd/rd dLGN units (**Fig 3G-H)**, with the latter showing strong ON bias (median ON–OFF bias index = 0.32 for Grm6, 0.69 for Brn3C, U = -7.87, p < 0.001, **Fig 3I**). However, the genotype difference in response persistence was not found in the dLGN, with both genotypes showing more transient responses in the brain (median transience index = 0.17 for both genotypes, U = -0.64, p = 0.520, **Fig 3J**).

ReaChR Brn3c rd/rd retinas showed less diversity in response polarity and transience compared not only to ReaChR Grm6 rd/rd, but also to published reports of the intact retina ^32,33^. This implies that ReaChR-driven activation is unable to recreate native visual response properties of many retinal ganglion cells (e.g. those with highly transient and OFF or ON/OFF polarity). To address this possibility more directly, we carried out a series of MEA recordings in visually intact retinas from mice containing ReaChR expression under the Brn3c promoter that are also heterozygous for Pde6b rd1 (termed ReaChR Brn3c rd het here; **Fig 4A**). In these animals we were able to record both the native photoreceptor response (at light intensity below ReaChR threshold, 13.95 log photons/cm^2^/s), and the isolated ReaChR responses (at high irradiance, 15.95 log photons/cm^2^/s, following pharmacological blockade of rod and cone signalling). This allowed us to perform a within-unit comparison of the photoreceptor-driven (PRC only) and ReaChR-driven (ReaChR only) responses (see methods for details, **Fig 4B**).

**Figure 4.**
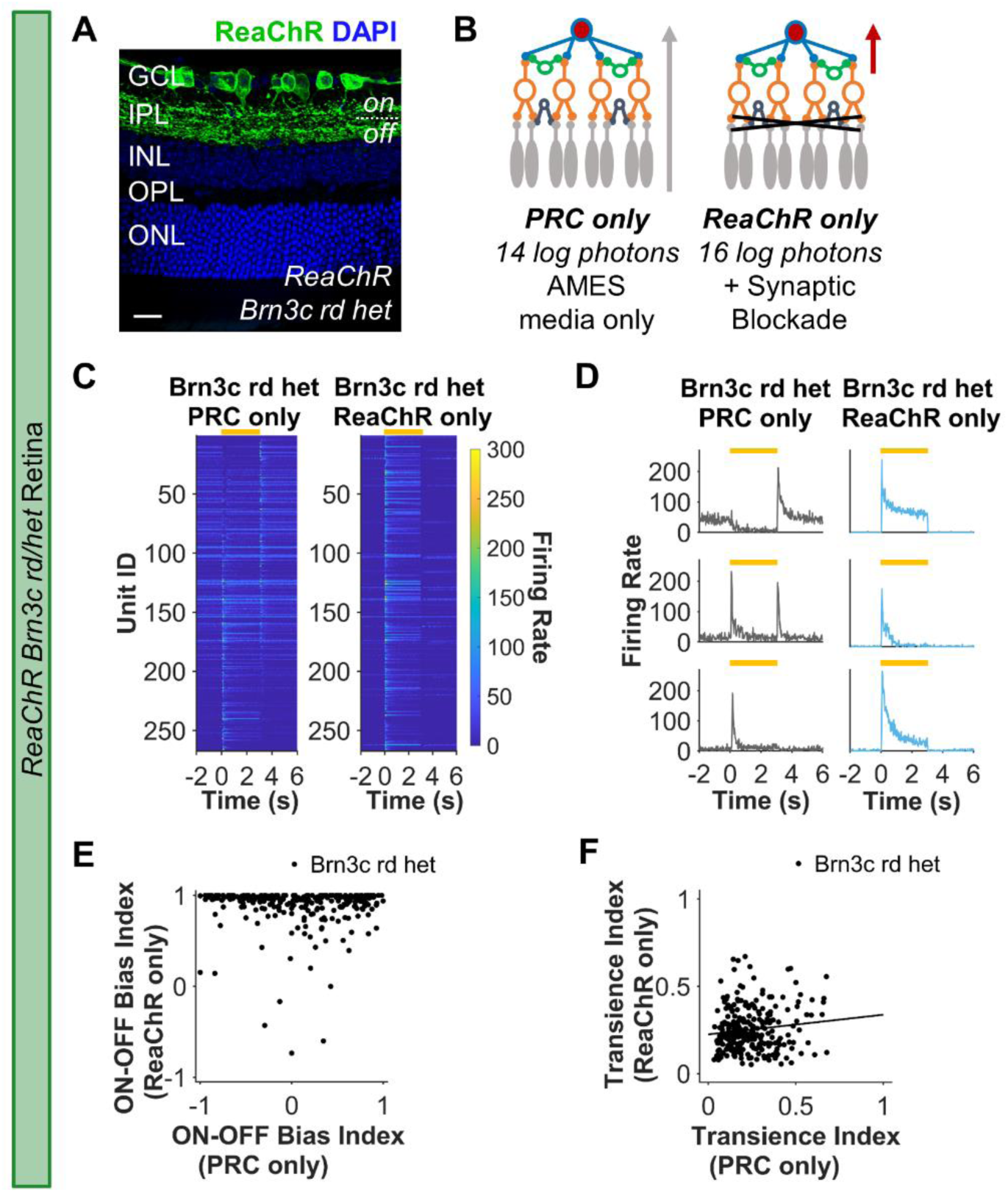
a) Immunohistochemistry from retinal sections from ReaChR Brn3C rd/het stained for DAPI (blue) and ReaChR-mCitrine (GFP). Scale bar = 20µm. b) Brn3c-positive RGCs are identified from ReaChR Brn3c rdhet recordings based on LR after synaptic blockade of rod/cone input. The photoreceptor-driven (PRC-only) responses recorded at lower light intensity, below threshold for ReaChR activation, in AMES media were compared to ReaChR-driven responses recorded at bright intensities in AMES media containing rod/cone blockers. c) Heatmap of mean PSTH for step stimulus (ON from 0-3s) for LR units ordered by bias index from OFF (top) to ON (high) in PRC-only condition. Each row represents the same individual unit recorded under PRC-only and ReaChR-only conditions. d) Example responses to step stimulus for 3 individual retinal units recorded under PRC- and ReaChR-only conditions. e) ON-OFF bias index f) Transience index Data in c-e) are from 267 units from ReaChR Brn3c rdhet (must be LR under both PRC and ReaChR-only conditions to be included). Yellow bar in c,d) shows timing of step stimulus

An overview of step responses suggested that across the entire population there was indeed more diversity in PRC-only compared to ReaChR response (**Fig 4C**). Moreover, we identified many units that either switched polarity, lost OFF response components or showed more sustained responses when activated via ReaChR compared to native photoreceptor input (**Fig 4C-D**). These could represent examples of units whose visual response properties are altered by ReaChR expression, but native ganglion cell responses can also change as a function of background light intensity ^43^.

Two lines of argument support the hypothesis that at least some of the differences in response properties between PRC and ReaChR conditions reflect a genuine disconnect between the native response of ganglion cells and that produced by direct optogenetic stimulation. Firstly, a unit-by-unit comparison confirms that a great deal of diversity in response polarity in the PRC condition is lost in ReaChR condition (**Fig 4D-E**), with ReaChR-only responses showing very strong ON bias. Secondly, there was no significant relationship in ON-OFF bias index of individual units under the two conditions (Pearson R = -0.02, p = 0.648, **Fig 4E**), which could be well-described by a flat line. Nevertheless, to more directly determine whether a reduction in response diversity is expected at high irradiance we recorded responses from wild type retinas under the same two intensities (13.95 and 15.95 log photons/cm^2^/s). These recordings showed equivalent diversity in ON-OFF bias at high vs low irradiance and strong correlation in this parameter at single unit level (**Supplementary Fig 1A-B,** Pearson R = 0.69, p < 0.001). Overall, these data confirm that at least some units that would be expected to have marked OFF response components instead become strongly ON biased under ReaChR stimulation.

ReaChR expression in RGCs likely contributed to the bias towards sustained responses seen in ReaChR Brn3c rd/rd, because there was weaker correlation in transience index between ReaChR vs PRC conditions (**Fig 4F**, Pearson R = 0.12, p = 0.038) than high vs low irradiance in wildtype (**Supplementary Fig 1C**, Pearson R = 0.42, p < 0.001). Interestingly, however, a degree of inter-unit variation in transience was retained in the ReaChR condition and this was weakly correlated with transience in the PRC condition (Pearson R = 0.12, p = 0.038). These latter findings suggest that transience may be partly an intrinsic property of RGCs irrespective of whether they receive visual input from photoreceptors or ReaChR.

### High Contrast Sensitivity Units Lacking in Brn3c RGC-targeting

We next examined how well each of the target cell types for optogenetic therapy recreated diversity in contrast sensitivity. In the dLGN, we saw a range of responses to the contrast chirp in both ReaChR Brn3c rd/rd and ReaChR Grm6 rd/rd animals (**Fig 5A**). This included examples of units showing graded increases in response across the contrast range; saturating responses at intermediate contrast; or responses to only the highest contrasts presented (**Fig 5B**). Extracting C50 from Naka-Rushton fits to contrast response functions revealed that the ReaChR Grm6 rd/rd retinal units were biased to slightly higher contrast sensitivity compared to ReaChR Brn3c rd/rd (median C50 = 61% for Grm6 and 69% for Brn3c, U = -3.20, p = 0.001, **Fig 5C**). The variation in contrast response was even more apparent in the retina (**Fig 5D**) with some units demonstrating large amplitude responses even at relatively low contrasts (**Fig 5E**). Comparison of C50 values revealed that such high contrast sensitivity was a particular property of the ReaChR Grm6 rd/rd retina (**Fig5F**) and accordingly, ReaChR Brn3c rd/rd units on average had significantly reduced contrast sensitivity (median C50 = 45% for Grm6 and 67% for Brn3c, U = -11.13, p < 0.001).

**Figure 5.**
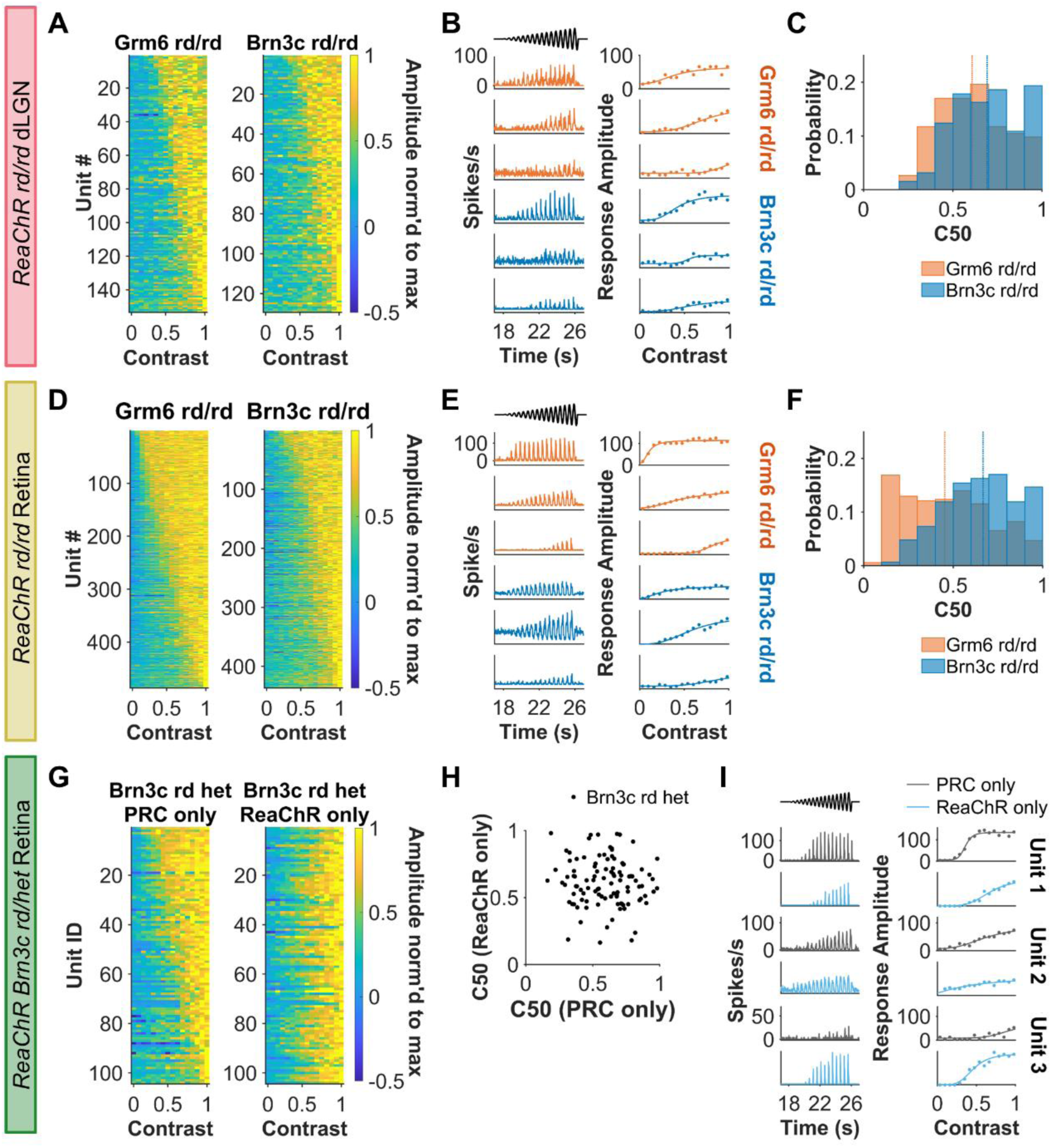
a,d,g) Heatmap of maximum normalised response amplitude across contrasts for LR units ordered by C50 from most (top) to least sensitive (bottom). Each row represents an individual unit and in g) each row represents the same unit recorded under different conditions (ranked based on C50 for PRC-only). b,e,i) Example firing rate to contrast chirp (left column) and contrast sensitivity function (right column) for 3 representative units. Data shown in h) are grouped by rows to show same unit under PRC- and ReaChR-only conditions. Timing of contrast chirp stimulus shown in black. c,f,h) Half maximal contrast (C50) derived from best-fit contrast response function. Scatterplot in h) shows C50 values derived from same unit under different conditions. Data in a-c) are from dLGN units (N = 153 for ReaChR Grm6 rd/rd and 129 for ReaChR Brn3C rd/rd), d-f) from rd/rd retina (N = 485 for ReaChR Grm6 rd/rd and 436 for ReaChR Brn3C rd/rd), and g-i) from ReaChR Brn3c rd het retina (N = 104 units)Contrast function must have R^2^ > 0.5 and LR units must have spiking in > 10% bins during contrast chirp to be included.

Brn3c retinas appear to lack high contrast sensitive units– but does this indicate a fundamental inability of direct optogenetic RGC activation to produce high contrast sensitivity? To address this question, we turned to comparing PRC and ReaChR only responses in visually intact ReaChR Brn3c rdhet animals (see above). This revealed that high contrast sensitivity (C50<0.5) units were rare under both PRC and ReaChR conditions (**Fig 5H**). The most straightforward explanation then is that high contrast sensitivity is rare in Brn3c positive ganglion cells. Further to the conclusion that ReaChR itself does not introduce a bias in contrast sensitivity, there was a spread of C50 values >0.5 in both conditions (**Fig 5H**) and it was possible to identify units showing both increased and decreased contrast sensitivity in ReaChR vs PRC conditions (**Fig 5I**). Turning to the question of whether ReaChR was able to recapitulate native contrast sensitivity of individual units, there was no significant correlation between C50 under ReaChR and PRC conditions in ReaChR Brn3c rdhet retinas (Pearson R = -0.02, p = 0.800, **Fig 5H**). However, analysis of this parameter in wild type retinas under the two irradiances used for ReaChR-only and PRC-only conditions revealed only a weak correlation (Pearson R = 0.15, p = 0.01, **Supplementary Fig 2**). It seems then that under these conditions C50 of individual units shows substantial plasticity over changes in irradiance, making it impossible for us to determine the extent to which the ReaChR-driven response recapitulates native response properties at single unit level.

### Temporal frequency tuning is more stereotyped with RGC targeting

We found dLGN units responded across the frequency range (1 to 8Hz) of the temporal chirp in both ReaChR Brn3C rd/rd and ReaChR Grm6 rd/rd (**Fig 6A**). This encompassed units with broad and narrow temporal frequency tuning (**Fig 6B**). The preferred temporal frequency (Peak TF) for each unit was extracted by fitting a Half-Gaussians function to average response amplitude at each temporal frequency. Peak TF was slightly higher across ReaChR Brn3C rd/rd than ReaChR Brn3c rd/rd units (median Peak TF = 1.72Hz for Grm6, 2.16Hz for Brn3c, U = -7.16, p < 0.001, **Fig 6C**). A more notable effect was a reduction in diversity in this parameter in ReaChR Brn3C rd/rd (**Fig 6D**, D = 0.33, p < 0.001, Kolmogarov-Smirnov test). This revealed that in the dLGN, ReaChR Grm6 rd/rd units had more variety in their temporal frequency tuning profiles (**Fig 6A**), including low-pass and bandpass units (**Fig 6B**), while the Brn3C dLGN which was dominated by units with bandpass tuning with peak TF at 2Hz.

**Figure 6.**
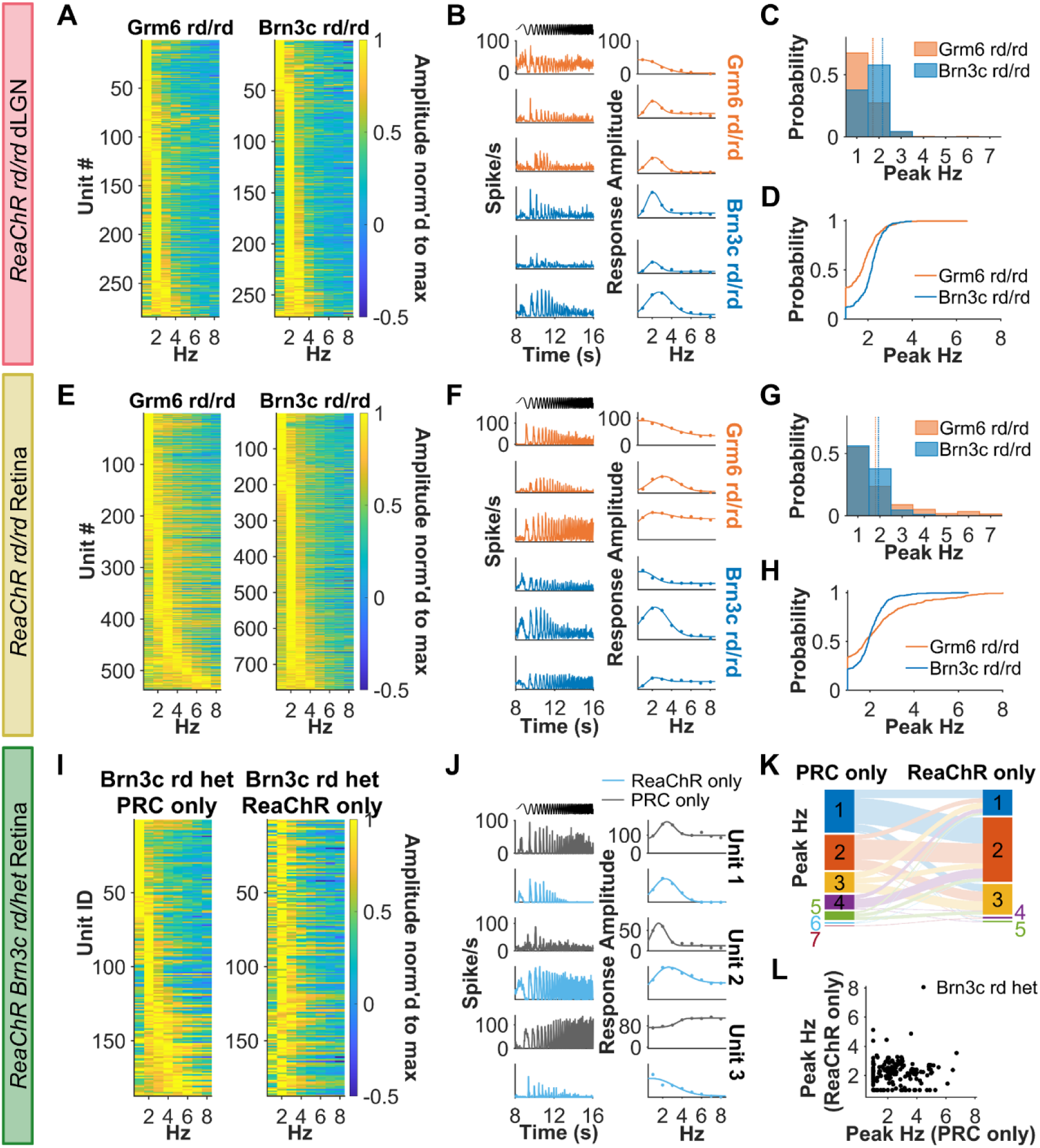
a,e,i) Heatmap of maximum normalised response amplitude across average frequency (Hz) for LR units ordered by peak temporal frequency (TF) from low (top) to high frequency (bottom). Each row represents an individual unit and in i) each row represents the same unit recorded under different conditions (ranked based on peak TF for PRC-only). b,f,j) Example firing rate to contrast chirp (left column) and contrast sensitivity function (right column) for 3 representative units. Data shown in j) are grouped by rows to show same unit under PRC- and ReaChR-only conditions. Timing of temporal chirp stimulus shown in black. c,g) Histogram and d,h) Cumulative distribution function for Peak TF (top) derived from best-fit temporal tuning function. k) Sankey diagram showing peak TF of units for PRC- and ReaChR-only conditions. Scatterplot in L) shows Peak TF values derived from same ReaChR Brn3c rdhet retinal unit under different conditions. Data in a-d) are from dLGN units (N = 285 for ReaChR Grm6 rd/rd and 270 for ReaChR Brn3C rd/rd), e-h) from retina (N = 285 for ReaChR Grm6 rd/rd and 270 for ReaChR Brn3c rd/rd) and i-l) from Brn3c rd het retina (N = 187 units). Best-fit temporal tuning profiles must have R2 > 0.5, LR units must have spiking in > 10% bins for contrast chirp, Gaussian spread > 0.51 (defined by at least 3 points) to be included.

Temporal frequency tuning was more diverse in the retina than the dLGN (**Fig 6E**). We found individual ReaChR Grm6 rd/rd retinal units (**Fig 6F**) with low-pass tuning, bandpass tuning, or strong responses at all temporal frequencies. In comparison, ReaChR Brn3c rd/rd units were more biased to 2Hz band-pass tuning, as in dLGN. Thus, although there was no significant difference in calculated Peak TF between genotypes (median peak TF = 1.79 for Grm6, 1.93 for Brn3c, U = -0.26, p = 0.796, **Fig 6G**), there was a difference in cumulative distribution - with ReaChR Brn3c rd/rd units more biased to 2Hz (**Fig 6H**, D = 0.16, p < 0.001, Kolmogarov-Smirnov test).

Examination of ReaChR Brn3c rdhet responses suggests that the 2Hz bias is a property of ReaChR in retinal ganglion cells. Thus, RGCs with diverse tuning profiles during photoreceptor-driven conditions realign towards 2Hz bandpass tuning under ReaChR-driven conditions (**Fig 6I-L**) to leave no statistically significant relationship between Peak TF under ReaChR vs PRC conditions (**Fig 6I**, Pearson R = -0.01, p = 0.918). Conversely, no such realignment was apparent in wildtype retinas under the same two light intensities (**Supplementary Fig 3,** Pearson R = 0.26, p < 0.001).

### RGC targeting produces an impoverished visual code

Having examined how targeting BC vs RGCs affects individual visual response properties we turned to a more holistic examination of the visual code. Applying community detection, we aimed to group units based on their response to the entire chirp stimulus as an approach to identifying information channels, defined by their response across different stimulus features. By pooling the ReaChR Brn3c and Grm6 rd/rd data with that from wildtype (photoreceptor-driven) retinas and then performing clustering and community detection, we hope to determine whether the optogenetic interventions: 1) differed in the number of parallel information channels recreated; 2) introduced bias in how units were distributed across these channels; and 3) closely recreated feature selectivity combinations found in wildtype mice.

Starting with the retina, 9 communities across 3 genotypes were identified (**Fig 7A-C**). ReaChR Grm6 rd/rd and wildtype units were distributed across all 9 communities, consistent with our previous report that ReaChR expression in ON bipolar cells can recapitulate much of the diversity of the wildtype visual code ^34^. By contrast, ReaChR Brn3c rd/rd units appeared in fewer clusters, being primarily restricted to communities 6, 7 and 8 (characterised by sustained ON responses, bandpass temporal tuning and intermediate contrast sensitivity). Only 1 Brn3c unit was found in community 3 (Off units with high contrast sensitivity and minimal temporal tuning) and no Brn3c units in community 4 (highly transient ON responses with high-pass temporal tuning and intermediate contrast sensitivity). The implication is that the method of optogenetic targeting had impacted the visual code, and indeed a shuffle test revealed that the distribution of units across communities was significantly different between ReaChR Grm6 and Brn3c rd/rd retinas (p = 0.001). While there was a small but significant difference in the distribution of units from WT and ReaChR Grm6 rd/rd retinas across communities (p = 0.025, as previously reported ^34^, the differences from wildtype were more stark in the ReaChR Brn3c rd/rd retina (p < 0.001), with Brn3c units mostly found mostly in communities (6-8) sparsely represented in wildtype retina.

**Figure 7.**
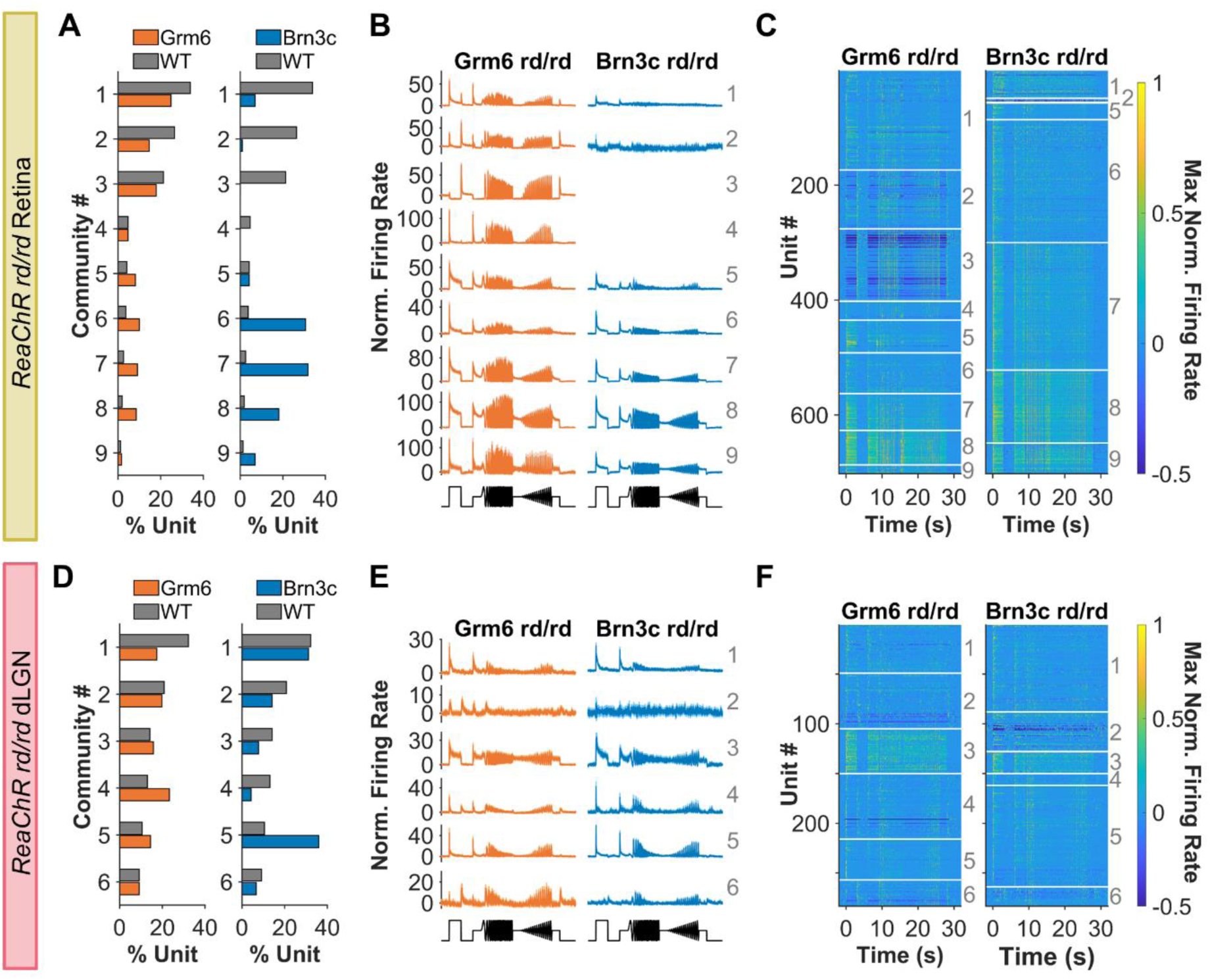
a,d) Distribution of a) retina (N = 702 units per genotype) and d) dLGN units (N = 283 units per genotype) across communities – downsampled to get same number in each genotype. b,e) Mean firing rate for units from each community (bold numbers) in b) retina and e) dLGN c,f) Heat map showing mean PSTH for individual units (each row) in each community. Boundaries of each community are shown with white lines and community identity is shown by bold number on right for c) retina and f) dLGN from rd/rd mice. Data from ReaChR Grm6 rd/rd mice in orange, ReaChR Brn3c rd/rd in blue and wildtype mice in grey. Labels for different communities are shown in grey text on b-c, e-f.

A similar pattern was observed in the dLGN, where we found 6 communities (**Fig 7D-F**). As in the retina, the distribution of units across these communities was significantly different between genotypes (p < 0.005) – with ReaChR Brn3c rd/rd units concentrated in communities 1 and 5 (transient ON with bandpass tuning and intermediate contrast sensitivity), while ReaChR Grm6 rd/rd units were more evenly distributed across the different response categories, suggesting that the more diverse visual code produced by ON BC targeting persists at higher levels of visual projection. As in the retina, the ReaChR Grm6 rd/rd responses more reliably recreated the visual code seen in WT animals, with similar distribution of units across communities (p = 0.154, shuffle test), while this difference between WT and Brn3c responses was more exaggerated (p = 0.009, shuffle test).

## DISCUSSION

Our aim was to provide a close comparison of therapeutic efficacy for retinal ganglion vs ON bipolar cell expression in optogenetic vision restoration. To this end we compared visual responses in transgenic mouse lines expressing the same optogenetic actuator (ReaChR) under the same promoter in targeted cell types across the retina, minimising the variation in extent and density of expression that is a feature of viral gene delivery methods. Our data confirm that ON bipolar cells provide clear advantages in terms of the quality of restored visual code. ON bipolar cell targeting did not impair sensitivity as previously reported and although response latency was increased compared to RGC targeting, it remained within the range for intact vision. ON bipolar cell targeting produced higher trial-to-trial reproducibility, but its most significant advantage lay in the richness of the restored visual code. Whereas ReaChR Brn3c rd/rd units converged to a relatively stereotyped response profile in terms of polarity, transience, temporal frequency tuning and contrast sensitivity, all of these characteristics showed greater variability in ReaChR Grm6 rd/rd. As a result, the ReaChR transgenic line with ON bipolar cell targeting better approached the diversity of visual information channels in the intact retina.

Some aspects of the ON BC advantage are consistent with *a priori* expectations based upon known retinal physiology. Visual signals introduced to ON bipolar cells must traverse the retina in order to reach ganglion cells and the brain. In doing so they may benefit from the information processing capacity of the inner retina in a way that visual signals introduced at the ganglion cell level cannot. The most obvious impact of such processing is the appearance of OFF and ON/OFF responses in ReaChR Grm6 rd/rd mice. The primary ReaChR response (cation conductance) should be the same in both ON bipolar and RGCs, but while the resultant light-dependent depolarisation can only appear as an ON excitation when introduced to RGCs, it can be transformed to OFF responses downstream from ON bipolar cells thanks to crossover inhibition with the retinal OFF pathway via AII amacrine circuitry ^44–50^. Similarly, temporal frequency tuning is known to be influenced both by intrinsic properties of ON BCs ^51^ and circuit mechanisms in the inner plexiform layer ^52–54^, providing plausible explanations for enhanced diversity in this characteristic in ReaChR Grm6 rd/rd mice. Diversity in response transience is also a property of neurons upstream of RGCs ^55^ and our data are consistent with the view that introducing ReaChR in ON bipolar cells allows more of that variability in response persistence to be recovered. Interestingly, however, the retention of a weak but significant correlation between this property in PRC and ReaChR-driven responses in the intact ReaChR Brn3c retina indicates that transience is defined to some extent in RGCs in such a way as to be accessible to direct optogenetic activation.

The more stereotyped responses of ReaChR Brn3c rd/rd imply a problem for the visual code in this genotype beyond its reduced diversity. Comparison of ReaChR and native photoreceptor-driven responses in visually intact rd het mice reveal that in many cases fundamental sensory response properties of units are different under direct optogenetic activation. Many OFF units switch to ON polarity and there are quantitative changes in transience, contrast sensitivity and temporal frequency tuning. In this way, firing patterns at an individual unit level convey quite different information about the visual scene under ReaChR activation. The extent to which this presents a problem for downstream processing remains uncertain and it is worth highlighting that some visual properties, such as response polarity, show substantial natural plasticity ^43,56,57^. That plasticity is particularly reported across changes in irradiance, and we have here included control recordings comparing response property changes within units between irradiances in wildtype mice. Shifts in most response parameters, including response polarity, transience and temporal tuning, were more common when switching from photoreceptor to optogenetic activation than when adjusting irradiance, confirming that the normal visual code is disrupted by ReaChR expression in RGCs beyond what is expected for natural plasticity. The exception is contrast sensitivity, which also shows large changes at a single unit level across irradiances, making it impossible to determine whether ReaChR in RGCs imposes additional alterations to diversity in this parameter.

Another theoretical advantage of ON bipolar cell targeting is that it may provide access to all retinal output channels in a way that may be hard to achieve for RGC targeting given cell-type specificity of viral transduction efficiency ^22^. The Brn3c Cre line provides an example of this challenge as it targets the subset of RGCs that are Brn3c-positive ^40^, comprising 27 of the 42 RGC functional subtypes identified in ^33^. Our parallel recordings of ReaChR and photoreceptor-derived responses in rd het mice provide an insight into the degree to which the incomplete coverage of RGC types contributes to the reduced visual code diversity in ReaChR Brn3c rd/rd. We find that the Brn3c positive cells show diversity in response polarity, transience, and temporal frequency tuning under photoreceptor driven conditions, confirming that the reduced diversity in these properties cannot be attributed solely to limitations in ganglion cell coverage. However, high contrast sensitivity cells do appear missing from the Brn3c population suggesting that incomplete coverage of the RGC types contributes to the reduced diversity in this property in ReaChR Brn3c rd/rd mice.

Although our data reveal several advantages of ON bipolar cell targeting, it is important to recognise that they do also provide encouragement for ganglion cell targeting. There may be good practical reasons for targeting ganglion cells in clinical practice including difficulties in efficiently transducing ON bipolar cells and the challenges of progressive inner retinal degeneration (although see ^34,58^). Our data extend the existing evidence that targeting ganglion cells can provide visual signals with helpful characteristics, such as a high spatiotemporal resolution ^18^. We found widespread and reproducible visual responses in ReaChR Brn3c mice across a range of irradiances, contrasts, and temporal frequencies. ReaChR Brn3c rd/rd also retain some diversity in response properties. Although strongly biased towards ON responses, there are a few rare examples of OFF excitation in the dLGN recordings. More encouragingly, single units in both retina and dLGN show surprising diversity in other visual responses parameters. Previous reports have seen a wide range of response transience following RGC-biased viral delivery ^16,18,35^ and we confirm that this is the case also when ReaChR expression is restricted to RGCs ^21^. We further show variability in contrast sensitivity, as well as two types of temporal frequency tuning profiles during RGC optogenetic targeting (low-pass and 2Hz bandpass). These findings highlight an interesting question of the extent to which, in normal vision, such properties are defined by intrinsic properties of individual RGCs versus the upstream circuit. Indeed, intrinsic mechanisms shaping visual feature extraction, such as contrast sensitivity, are beginning to be described ^59^. From a practical perspective, these properties contribute to a richer visual code following RGC targeting than may have been initially anticipated.

An important property shared by ON bipolar and ganglion cell interventions was sensitivity normalisation. Natural photoreceptors face the challenge of encoding small local modulations in light intensity across big differences in background light with a variety of adaptation mechanisms. A reasonable concern with optogenetic interventions is that, absent such mechanisms, restored vision would have a narrow sensitivity range, saturating at bright backgrounds. In fact, we find evidence of sensitivity normalisation in both ReaChR Grm6 rd/rd and ReaChR Brn3c rd/rd, with contrast sensitivity retained across brighter irradiances. Inner-retinal mechanisms of light adaptation may contribute to this ability in ReaChR Grm6 rd/rd ^60–64^, but the ReaChR Brn3c rd/rd data implies that it may also be an intrinsic property of ReaChR activity and/or host cell physiology.

The decision of which cell types to target for optogenetic therapy in clinical application will be a complex one and may encompass patient-specific considerations such as inner retinal integrity, as well as practical challenges of viral gene delivery ^65^. Our side-by-side comparison of the quality of restored vision from ON bipolar vs RGC targeting suggests that the theoretical advantages of introducing visual signals as early as possible to the circuit are indeed apparent in a richer visual code. However, they also confirm an impressive ability to encode dynamic visual stimuli across a range of irradiances with either targeting method and thus provide general encouragement for this therapeutic avenue.

## MATERIALS AND METHODS

This study includes some electrophysiology data from units in ReaChR Grm6 rd/rd and visually intact mice previously analysed in Rodgers et al ^34^. For retinal recordings, this original dataset has not been altered, while for dLGN dataset, additional recordings from ReaChR Grm6 rd/rd mice have been added. All data from ReaChR Brn3c rd het or rd/rd mice was newly generated for this study and has not been previously published. Existing data from ReaChR Grm6 rd/rd and WT mice were analysed, alongside new recordings from ReaChR Brn3c animals, using new criteria to identify light responsive units.

### Animals

MEA recordings were from: 5 retinas from 5 *ReaChR Grm6^Cre/WT^ Pde6b^rd1/rd1^* mice; 7 retinas from 4 *ReaChR Brn3c^Cre/WT^ Pde6b^rd1/^*; 5 retinas from 4 *ReaChR Brn3c^Cre/WT^ Pde6b^rd1/WT^* mice. MEA recordings from visually intact animals were from: 3 retinas from 3 *ReaChR Grm6^WT/WT^ Pde6b^rd1/WT^* mice; 9 retinas from 8 *C57Bl/6* mice (Envigo) as described in Rodgers et al ^34^. LGN recordings were from: 16 placements from 7 *ReaChR Grm6^Cre/WT^ Pde6b^rd1/rd1^* mice 14 placements from 6 mice from *ReaChR Brn3c^Cre/WT^ Pde6b^rd1/rd1^*mice LGN recordings from visually intact animals were from 11 placements from 4 *C57Bl/6* mice (University of Manchester,) as described in Rodgers et al ^34^. *ReaChR Grm6^Cre^ Pde6b^rd1^* mice were produced at University of Oxford, *ReaChR Brn3c^Cre^ Pde6b^rd1^* mice were produced at University of Manchester. All animals were given water and food *ad libitum*, kept under 12:12 light-dark cycle and group housed. Home cage lighting intensity was below threshold for ReaChR activation. All experiments were conducted in accordance with the Animals Scientific Procedures Act of 1986 (United Kingdom) and approved by ethical review committees at University of Oxford and University of Manchester.

### Transgenic mice

This study uses the ReaChR Grm6 rd strain, previously described in Rodgers et al.^34^, created by breeding Grm6^Cre/WT^ (MGI:4411993, kindly shared by Robert Duvoisin, Oregon Health and Science University, USA) with ReaChR-mCitrine mice (MGI: 5605725) obtained from Jax laboratories (#026294). ReaChR Grm6 rd/rd mice were bred to be homozygous for ReaChR-mCitrine, heterozygous for Grm6 Cre and homozygous for Pde6b rd1. We also produced a new transgenic line – ReaChR Brn3c rd. Brn3c^Cre/WT^ mice (MGI: 7470766, kindly shared by Tudor Badea, National Eye Institute, NIH,USA ^40^) were bred with homozygous *ReaChR Grm6^WT/WT^ Pde6b^rd1/rd1^*from the *ReaChR Grm6 rd* colony maintained at the University of Manchester. Mice were bred to be homozygous for ReaChR, heterozygous for Brn3c Cre and either heterozygous (ReaChR Brn3c rd het) or homozygous (ReaChR Brn3c rd/rd) for *Pde6b^rd1^*. These mice express floxed *ReaChR-mCitrine* transgene in Brn3c-positive RGCs which express Cre recombinase. Genotyping was performed using primers to amplify Brn3c Cre (Fwd = 5’-CCGGGGTATAAATGCTGTGG, Rev = 5’-CCTCATCACTCGTTGCATCG,

411bp band), ReaChR and rd1 alleles (as described in Rodgers et al ^34^) or using an external genotyping service (Transnetyx). At 5 months old, *Pde6b^rd1/WT^* mice are retinally degenerate, with complete loss of rods and cone photoreceptors. *Pde6b^rd1/WT^* are visually intact and possess rod and cone photoreceptors. Unless stated otherwise, mice are on a mixed C57Bl/6 x C3H background.

### Immunohistochemistry

Retinal sections were immunostained as previously described ^23,66^. mCitrine tag of ReaChR-mCitrine transgene was labelled using chicken anti-GFP polyclonal primary antibody (1:1000, GFP-1020, AVES labs). Donkey anti-chicken 488 (1:250, T03-545-155, Jackson Immunoresearch) was used as secondary antibody. Fluorescence images were acquired using inverted LSM 710 laser scanning confocal microscope (Zeiss) with Zen 2009 image acquisition software (Zeiss). Individual channels were collected sequentially. Excitation laser lines were 405nm and 488nm with emission at 440-480 and 505-550 respectively. Z-stack was acquired using a x40 objective, with images collected every 1µm in z-axis. Global enhancement of brightness and contrast were applied to maximum intensity projection using ZenLite 2011 software (Zeiss).

### In vivo electrophysiology

Mice were anesthetised using urethane (intraperitoneal injection, 1.4-1.5g/kg) and placed in stereotaxic frame. An incision was made through scalp to expose surface of skull. A small hole was drilled 2.2mm lateral and 2.2mm posterior from bregma. The contralateral pupil was dilated using 1% atropine in saline (Sigma-Aldrich) and kept lubricated using mineral oil or Lubrithal (Dechra). Multi-electrode arrays (A4x16-Poly2-5mm-23s-200-177-A64, NeuroNexus) were coated in CM-DiI (Fisher Scientific); positioned at 2.2mm lateral and 2.2mm posterior relative to Bregma and inserted to depth of 2.5-3mm to target lateral geniculate nucleus, confirmed by presence of light responsive units to short steps of white light (2-5s ON, 10s OFF, 10 repeats. 16log photons/cm^2^/s). Once light responses were identified, mice were dark adapted for 20-30mins, allowing neuron activity to stabilise. Signals were acquired using Recorder 64 system (Plexon), amplified (x3000), high-pass filtered at 300Hz, digitised at 40kHz and stored in 16bit continuous format. Some additional recordings were performed by raising recording probe and moving 0.2mm posterior or anterior before re-inserting into the dLGN. After experiments were complete, mice were killed by cervical dislocation and brains were fixed in 4% paraformaldehyde. Single unit activity was isolated using Kilosort ^67^ and manually checked in Offline sorter (Plexon).

### Retinal MEA recordings

Mice were culled by cervical dislocation, enucleated and retinas dissected under dim red light. Retinas were positioned ganglion-cell side down in MEA chambers (Multi Channel Systems) containing 252 electrodes (30µm in diameter, spaced 100µm apart). MEAs were then inserted in MEA2100-256 recording system (Multi Channel Systems) and positioned in light path of inverted Olympus IX71 microscope. Retinas were kept at 34°C and were continuously perfused with AMES media gassed with 95% O2 and 5% CO2. Neural signals were collected, amplified and digitised at 25kHz using MCS Experimenter software (Multi Channel Systems). Retina were dark adapted for at least 30min before stimuli presentation to allow neural activity to stabilise. Single units were isolated from retinal MEA data using SpikeSorter software (Version 4.77b Nicholas Swindale, UBC). Raw data was filtered using a high pass 4-pole 500Hz Butterworth filter. Event detection was based on 4-5x median noise signal, with window width of 0.24ms. Automatic spike sorting results were manually checked using SpikeSorter software and Offline Sorter (Plexon).

For *ReaChR Brn3c rd het* retinal recordings, retinas were first perfused with AMES media during initial stimulus presentation, before being perfused AMES with synaptic blockers during second round of stimuli. The following pharmacological blockade was used to isolate ReaChR-driven responses in Brn3c-positive ganglion cells: 100 μm L(+)-2-amino-4-phosphonobutyrate (L-AP4) (group III metabotropic glutamate receptor agonist), 40 μm 6,7-dinitroquinoxaline-2,3-dione (DNQX) (AMPA/kainate receptor antagonist), and 30 μm d-2-amino-5-phosphonovalerate (d-AP5) (NMDA receptor antagonist, Tocris).

### Visual stimuli

For *in vivo* electrophysiology experiments, light was delivered using a CoolLED pE-4000 light source via a liquid light guide connected to a diffuser (Edmund Optics). This was positioned ∼5mm from eye contralateral to hemisphere containing recording site. White light was used for all stimuli (output from 4 LEDs at 385nm, 470nm, 550nm and 660nm). Neutral density filters were inserted in light path to produce intensity range from 12.99-16.97 effective log photons/cm^2^/s.

For retinal MEA experiments, light was delivered using a white LED light source with daylight spectrum (ThorLabs, SOLIS-3C), with stimuli generated by an arbitrary waveform generator (RS components, RSDG2000X series). Neutral density filters (ThorLabs) in a motorised filter wheel were used to control intensity of light stimuli from 11.9-17.4 log photons cm^2^/s. Devices were automatically controlled and synchronised by a Digidata 1440A digital I/O board (Axon Instruments, Molecular Devices, USA) and a PC running WinWCO software (J Dempster, Strathclyde University, UK).

Responses were recorded to full-field chirp stimuli consisting of 3s step from dark to 100% intensity, followed by 2s of dark, 2s at 50% intensity, 8s temporal chirp (accelerating sinusoidal modulation at 100% contrast from 1-8Hz at 1Hz/s), 2s at 50% intensity, 8s contrast chirp (2Hz sinusoidal modulation from 3% to 97% contrast), 2s at 50% intensity and 3s of dark. Chirp stimuli were presented from lowest to brightest intensity.

### Quantification and statistical analysis

Unless otherwise specified, graphs show mean with error bars showing standard error of the mean, sample size is given in figure legends and refers to number of retinal and LGN units. Comparisons between individual ReaChR Grm6 rd/rd and ReaChR Brn3c rd/rd units used Mann-Whitney U-tests, with significance determined as p < 0.05, at 15.95 log photons/cm^2^/s for LGN and 16.97 log photons/cm^2^/s for retina (unless stated otherwise). These irradiances were used as they produced responses closest to 75% maximum, providing a sub-saturating response with large sample size and good signal to noise ration.

#### Identifying light responsive units

Peristimulus time histograms (PSTH) with 25ms bin size were generated. Units with low spike firing rates (<10% of bins containing spiking activity) and spiking activity in <8 trials were excluded from further analysis. Light responsive (LR) units were identified using a shuffle test based on correlation across trials. A significance threshold of p < 0.0001 was used.

#### Response Amplitude and Irradiance Response Curves

Normalised firing rate was calculated by subtracting average baseline firing during 2s before onset of 3s step stimulus. Response amplitude was defined as max normalised firing rate during 3s step or 3s after step stimulus to capture both ON and OFF responses. The response amplitude of units that were LR to brightest intensity tested (16.97 log photons/cm^2^/s for retina and 17.4 log photons/cm^2^/s for LGN) was then averaged for each retina or LGN electrode placement and plotted against stimulus intensity. For retina, this data was fit with irradiance response curve using Hill-slope ^34^ with 4 free parameters (top, bottom, slope, EC50) to estimate photosensitivity.

#### Quality Index

To assess response reproducibility, PSTH with 200ms bin size was used to calculate to quality index ^32^, was generated for LR units at brightest intensity tested for each genotype.

#### Response Latency to Step onset

Latency to onset of step stimulus was based on PSTH with 25ms bin size and was calculated as timing of first bin to exceed 95% confidence limit in 1s after onset of step stimulus. This 95% confidence limit was based on 2 standard deviations of baseline firing during 1s before onset of light step. Units which did not exceed this threshold, such as OFF units, were excluded from this analysis.

#### Response Polarity and Transience

Units were grouped into response categories (ON transient, ON sustained, ON-OFF and OFF) using objective criteria, as described in Rodgers et al ^34^. ON-OFF bias index ^68^ was used to assess response polarity and was calculated as ratio of spike firing during 500ms after onset (ON_firing) and 500ms after offset (OFF_firing) of light step. This produces scale from -1 (firing to OFF only) to 0 (equal firing for ON and OFF) to 1 (firing to ON only).

Transience index ^35,68^ was used to test response persistence. PSTH with 25ms bin size was normalised to maximum firing rate during 3s step. Area under curve was then calculated for 1s after stimulus onset for ON units (defined as ON-OFF bias index > -0.33) or 1s after stimulus offset for OFF units (ON-OFF bias index < -0.33). This produces a scale from 0 (highly transient) to 1 (highly sustained with identical response across all bins tested).

#### Analysis of ReaChR Brn3c rdhet and WT retinas

In order to identify Brn3c positive units and compare their photoreceptor and ReaChR-driven activity, we identified units from ReaChR Brn3c rdhet retinas that were both 1) LR at light intensity below threshold for ReaChR activation (13.95 log photons/cm^2^/s) during perfusion with standard AMES *and* 2) LR at light intensity above threshold for ReaChR activation (15.95 log photons/cm^2^/s) during perfusion with AMES containing synaptic blockers. Activity under former condition was defined as photoreceptor-driven, while activity recorded under latter was defined as ReaChR-driven.

For analysis of wildtype retinas across intensities, we compared units that were LR at intensities above (13.95 and 15.95 log photons/cm^2^/s) during perfusion with standard AMES. As these mice do not possess ReaChR, responses are driven by rod and cone photoreceptors under both conditions.

#### Contrast sensitivity

To assess contrast sensitivity, we used PSTH with 25ms bin size. Response amplitude to each sinusoidal modulation was calculated as maximum firing – minimum firing rate during 0.5s each contrast was presented. These were then normalised to response amplitude during 0.5s before contrast chirp onset and plotted against Michelson contrast and fit with Naka-Rushton function ^69^ with 4 free parameters (top, bottom, slope and C50) using least squares minimisation. C50 was constrained between 0 and 1, and slope was constrained between 0 and 10. Only units with curve fits where R^2^ > 0.5 and spiking in >10% of bins were used for comparison of contrast sensitivity parameters.

#### Temporal frequency tuning

For temporal tuning, we used PSTH with 25ms bin size. The mean response amplitude (based on maximum firing-minimum firing during each sinusoidal modulation) was calculated for each temporal frequency. This data was fitted with a half-Gaussian model ^70^ with 5 free parameters (low baseline, high baseline, Gaussian spread, peak response amplitude and peak temporal frequency) using least squares minimisation. Peak temporal frequency was constrained between 1-8Hz. Only units with curve fits where R2 > 0.5, spiking in > 10% of bins and Gaussian spread > 0.51 were used for comparison of temporal frequency tuning parameters.

#### Community Detection

Sparse principal components (sPCs) were generated for 3 parts of chirp stimulus (Step at 0.5-4.5s, temporal chirp at 6.5 – 15.5s and contrast chirp at 17.5-24.5s) from PSTH with 50ms bin size using SPaSM toolbox ^71^. sPCs were generated based on pooled data from wildtype, ReaChR Grm6 rd/rd and ReaChR Brn3c rd/rd recordings. For retina, data was used from LR units at 15.95 (ReaChR Grm6 and Brn3c) and 12.95 photons/cm^2^/s (WT), while for LGN, data was used from LR units at 16.97 (ReaChR Grm6 and Brn3c) and 14.99 photons/cm^2^/s (WT). Units were randomly downsampled to match sample size of genotype with fewest units, n = 702 units for retina and n = 283 for LGN. After discarding sPCs accounting for <1% of variance, we extracted 57 sPCs for retina and 74 sPCs for dLGN with 5 non-zero timebins. sPCs were clustered using a Gaussian mixed model with random initialisation. Optimal number of clusters was determined by lowest Bayesian information criteria and Bayes factor < 6 (as in ^72^). Clustering was repeated 50 times and used to generate a pairwise similarity matrix. A community detection algorithm based on this similarity matrix, using Brain connectivity toolbox ^73^, was then used to group units into communities. Communities with <5 units were excluded from further analysis. Distribution of units across communities was compared between genotypes using a shuffle test, as described previously ^34^.

## DATA AVAILABILITY

Data reported in this paper will be shared by the lead contact upon request. Any additional information required to reanalyse the data reported in this paper is available from the lead contact upon request. The *ReaChR Brn3c rd* and *ReaChR Grm6 rd* transgenic mice are available subject to the completion of material transfer agreements.

## ACKNOWLEDGEMENTS

This work was funded by MRC grant (MR/S026266/1) awarded to MH, SH, SP and RL. ML was funded by grants from ProRetina Foundation (Pro-Re/Projekt/Gi-Wh-Li.04.2021) and German Research Foundation (LI 2846/6–1). AEA is funded by a Sir Henry Dale Fellowship, jointly funded by the Wellcome Trust and the Royal Society (Grant Number 218556/Z/19/Z). RS is funded by a Sir Henry Dale Fellowship, jointly funded by the Wellcome Trust and the Royal Society (Grant Number 220163/Z/20/Z).

## AUTHOR CONTRIBUTIONS

JR, SH, MH, and RJL designed the study. SH and JR performed experiments. TB provided Brn3c Cre mice. ML, AE, AEA, SNP and RS provided tools for data analysis, which was conducted by JR. JR, MW and RJL wrote the manuscript, which was edited and approved by all authors.

## SUPPLEMENTARY FIGURE LEGENDS

**Supplementary Figure 1.**
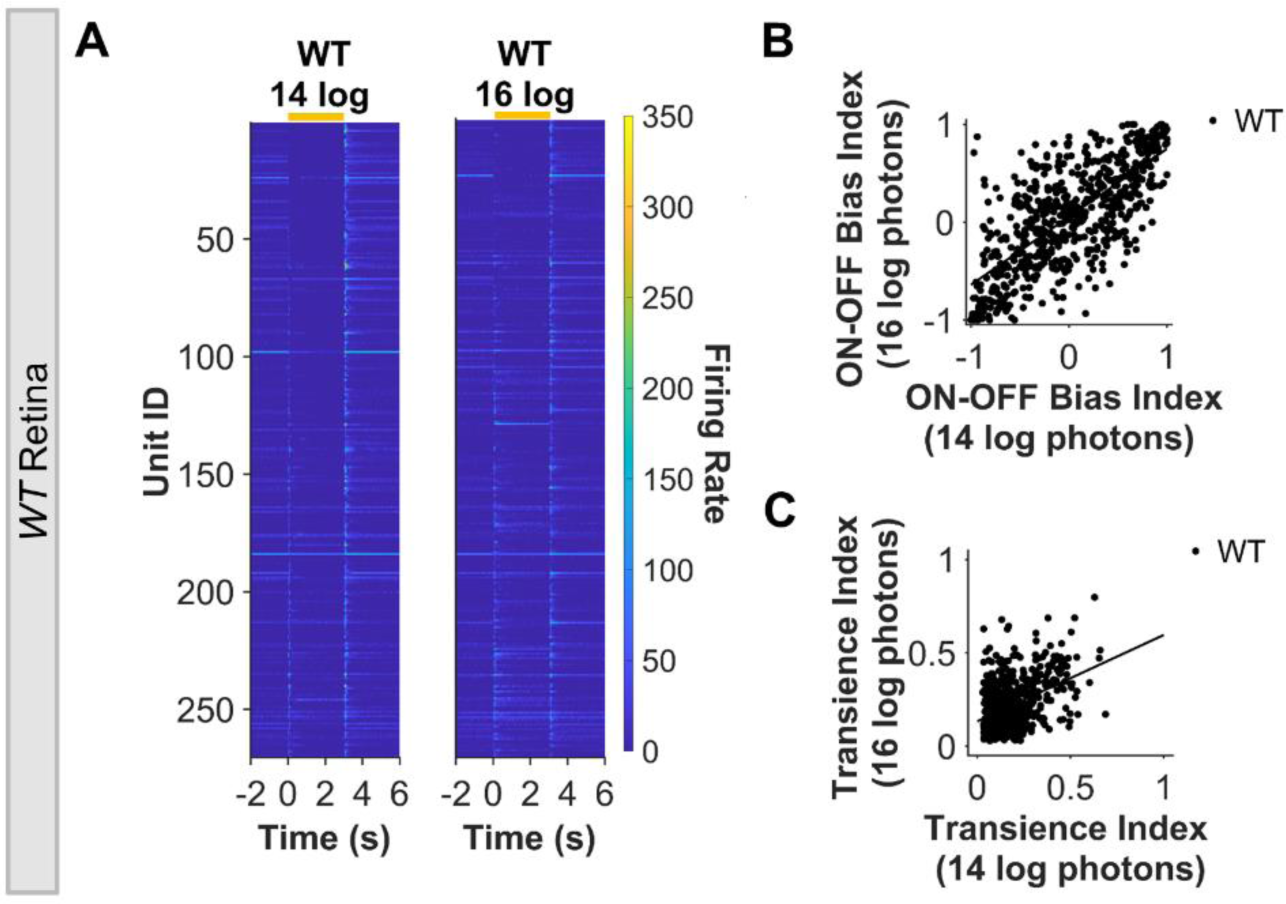
**a)** Heatmap of mean PSTH for step stimulus (ON from 0-3s) for LR units in WT retina ordered by bias index from OFF (top) to ON (high) under 13.95 and 15.95 log photons/cm2/s. Each row represents an individual unit and in two panels each row represents the same unit recorded under different conditions (ranked based on ON-OFF bias for 13.95 log photons). **b)** Scatterplot of ON-OFF bias index for same WT units at 13.95 and 15.95 log photons/cm2/s. **c)** Scatterplot of Transience index for WT units at 13.95 and 15.95 log photons/cm2/s. Data are from N = 630 WT retinal units (must be LR under both intensities to be included)

**Supplementary Figure 2.**
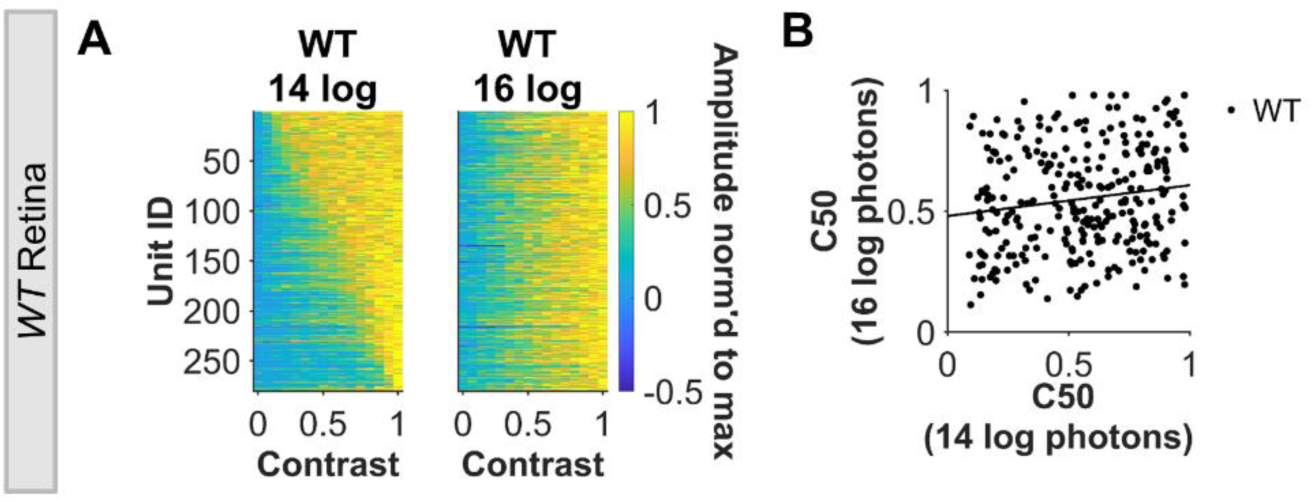
**a)** Heatmap of maximum normalised response amplitude across contrasts for LR WT retinal units. Each row represents an individual unit in two panels each row represents the same unit recorded under different conditions (ranked based on C50 for ND4). **b)** Scatterplot of C50 for same WT units at 13.95 and 15.95 log photons/cm2/s. Data are from N = 280 WT retinal units.

**Supplementary Figure 3.**
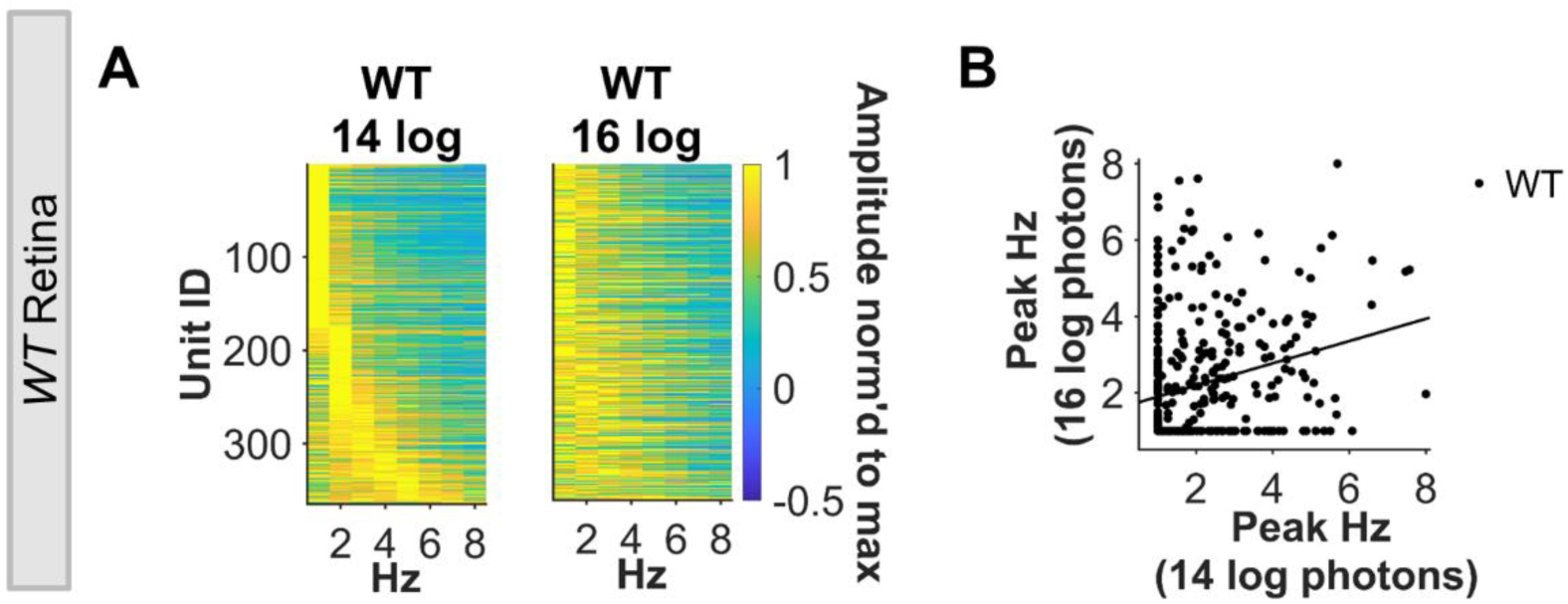
**a)** Heatmap of maximum normalised response amplitude across frequencies for LR WT retinal units. Each row represents an individual unit in two panels each row represents the same unit recorded under different conditions (ranked based on peak TF for ND4). **b)** Scatterplot of peak TF for same WT units at 13.95 and 15.95 log photons/cm2/s. Data are from N = 369 WT retinal units.

